# Dexrazoxane as a viable microsporidia control agent in *Anopheles gambiae*

**DOI:** 10.1101/2024.10.14.617874

**Authors:** Tiago G. Zeferino, Luís M. Silva

## Abstract

Microsporidia have long been proposed as biological agents for controlling disease vectors and the parasites they transmit. However, their study in vector biology has been constrained due to challenges in manipulating microsporidia within hosts. In this study, we investigated the effect of Dexrazoxane, a candidate drug against microsporidiosis, on the establishment and development of *Vavraia culicis* infection in its natural host, the mosquito *Anopheles gambiae*, the main malaria vector. Our findings show that Dexrazoxane significantly reduces spore load, particularly in mosquitoes reared individually, without affecting the overall infection success of the parasite. This result aligns with studies in *Caenorhabditis elegans*, where Dexrazoxane inhibited new spore production without hindering initial spore integration into the host gut cells. Dexrazoxane’s DNA topoisomerase II inhibitor mechanism likely explains its impact on mosquito development, as larvae exposed to the drug failed to emerge as adults. These findings highlight Dexrazoxane’s potential as a viable tool for controlling microsporidia in adult mosquitoes and hope to enhance the study of mosquito-microsporidia interactions. Further research is required to explore its broader application in vector-borne disease control, including malaria.

## Introduction

Microsporidia are common obligate parasites of many animals, and their prevalence has been increasingly reported over the past decades across a wide range of taxa, sparking a surge in microsporidia studies (Becnel and Weiss, 2014; Bojko et al., 2022; Franzen, 2008; Han et al., 2020; Seatamanoch et al., 2022; Wadi and Reinke, 2020; Weidner and Overstreet, 2021). While research in human-infecting microsporidia mainly focuses on clearing them (i.e., microsporidiosis) (Han et al., 2021), in invertebrates such as insects, their study is more nuanced, as evidence suggests microsporidia may serve as biological control agents for certain pests (Bjørnson and Oi, 2014) and vector species, and the diseases they transmit (Becnel and Andreadis, 2014; Bukhari et al., 2022; Koella et al., 2009).

Nearly half of the microsporidia species that infect insects have specialised in targeting dipterans, such as flies and mosquitoes, which are vectors for several human diseases. Among those infecting mosquitoes, *Vavraia culicis* (Silva et al., 2025; Vavra and Becnel, 2007) is one of the most well-studied microsporidia species (Agnew et al., 1999; Biron et al., 2005; Desjardins et al., 2015; Kelly et al., 1981; Rivero et al., 2007; Silva and Koella, 2024a, 2024b; Sy et al., 2014; Vyas-Patel, 2023; Zeferino and Koella, 2024). This generalist intracellular parasite is known to infect several mosquito genera, including *Anopheles gambiae*, the major vector of malaria (Zeferino and Koella, 2024). Infection typically occurs during the larval stage when mosquito larvae ingest spores along with food in water bodies. Once the spores penetrate the host’s gut wall, they develop and produce new spores.

The limited immune response of *An. gambiae* to this particular microsporidian species (Silva, 2024) allows for a steady progression of the infection, characterised by continuous consumption of host resources and accumulation of spores in the body cavity, leading to costs in host longevity (Agnew et al., 2004; Zeferino and Koella, 2024) and fecundity (Silva and Koella, 2024a).

Although *An. gambiae* incurs significant costs from *V. culicis* infection, it comes with a benefit in protection against co-infection with *Plasmodium spp*. by negatively affecting oocyst development (Bargielowski and Koella, 2009; Lorenz and Koella, 2011). These findings led the community to hypothesise that microsporidian parasites, such as the one in this study, could be used as biological control agents for mosquitoes and their parasites by reducing the mosquito lifespan and parasite transmission to mammal hosts, respectively (Bargielowski and Koella, 2009; Bukhari et al., 2022; Lorenz and Koella, 2011). However, the mechanisms underlying this interaction remain unclear, partly due to the technical challenges of manipulating microsporidian infections within mosquito hosts.

Hence, to facilitate the control and study of microsporidians within the mosquito host, a candidate chemical, Dexrazoxane, recently shown to have anti-microsporidian activity in *Caenorhabditis elegans* (Murareanu et al., 2022) was tested. Due to its antimitotic and redox activity, Dexrazoxane has been extensively used as a protector agent against the cardiotoxicity of chemotherapy in humans (Vejpongsa and Yeh, 2014). Its mechanism of action is similar to ethylenediaminetetraacetic acid (EDTA) but diffuses into cells more readily (Hasinoff et al., 1995), where it binds to iron and lowers oxidative stress formation (Eneh and Lekkala, 2020). However, its role against microsporidia has only recently been studied. For instance, in *C. elegans*, Dexrazoxane is known to inhibit *Nematocida parisii* development after it has established itself in the gut cells (Murareanu et al., 2022). While it does not prevent initial infection, it suppresses the production of new spores, thereby hindering disease progression and transmission. Given its promising effects on microsporidia infection in *C. elegans*, we aimed to test whether these effects extend to microsporidian infections in mosquitoes, using *An. gambiae* and *V. culicis* as the experimental system. Our findings suggest that Dexrazoxane’s effect might be applicable to more model systems than previously thought, extending to mosquito-microsporidia interactions. This result may enhance our understanding of microsporidia interactions with their host and other parasites and could potentially inform and fuel future vector-borne disease control strategies.

## Materials and Methods

### 1. Experimental system

We used the *An. gambiae s.s* Kisumu strain (Vulule et al., 1994), which had been maintained at our standard laboratory conditions (about 600 individuals per cage, constant access to 6% sucrose solution, 26 ± 1ºC, 70 ± 5% relative humidity and 12 h light/dark) for many years before the experiments.

The microsporidian *V. culicis floridensis* was provided by J.J Becnel (USDA, Gainesville, FL, USA). Originally discovered in *Ae. albopictus*, this parasite was later found to parasitise several mosquito genera, including *Aedes, Culex, Anopheles, Culiseta, Ochlerotatus* and *Orthopodomyia* (Andreadis, 2007; Fukuda et al., 1997; Kelly et al., 1981). We maintained it by alternately infecting *Aedes aegypti* and *An. gambiae* to ensure its status as a generalist parasite, as described previously (Zeferino and Koella, 2024).

*V. culicis* is an obligatory, intracellular parasite. Mosquitoes are infected when, as larvae, they ingest the spores along with their food. *V. culicis* culture and experiments described below used a dose of 10,000 spores per larva, as previous preliminary experiments have shown that it infects 100% of the mosquitoes at this dose. After several rounds of replication, the parasite produces new infectious spores that spread to the gut and fat body cells. The spores are released to larval sites when infected larvae die, adults die on the surface of water, or eggs covered with spores are laid onto the water’s surface.

### 2. Rearing and maintenance of experimental mosquitoes

Freshly hatched (0-3 hours old) *An. gambiae* larvae were individually placed into 12-well culture plates, each containing 3 ml of deionized water. The larvae received Tetramin Baby® fish food every day according to their age (0.04, 0.06, 0.08, 0.16, 0.32 and 0.6 mg/larva respectively on days 0, 1, 2, 3, 4 and 5 or older (Kulma et al., 2013)).

#### 2.1 Larvae exposure to Dexrazoxane

In the first experiment **(Fig. 1)**, we assessed the impact of Dexrazoxane (Cat# 05587, Sigma-Aldrich, St. Louis, MO, USA) on larval development and spore growth. Since the mosquito larvae were reared in deionized water with Dexrazoxane, we hypothesized that a long exposure time might differently affect the mosquito fitness traits. Therefore, we not only tested three different doses (0 µM, 30 µM, and 60 µM), but also tested for the impact of one administration at one of three time points (four, five, or six days after egg hatching) for infected and uninfected larvae, in a fully combinatorial design. The larvae’s survival and pupation were tracked daily. Since mosquitoes only have detectable sporulation in the adult stage (Silva and Koella, 2024a), spores were uncountable for this experiment.

**Figure 1.**
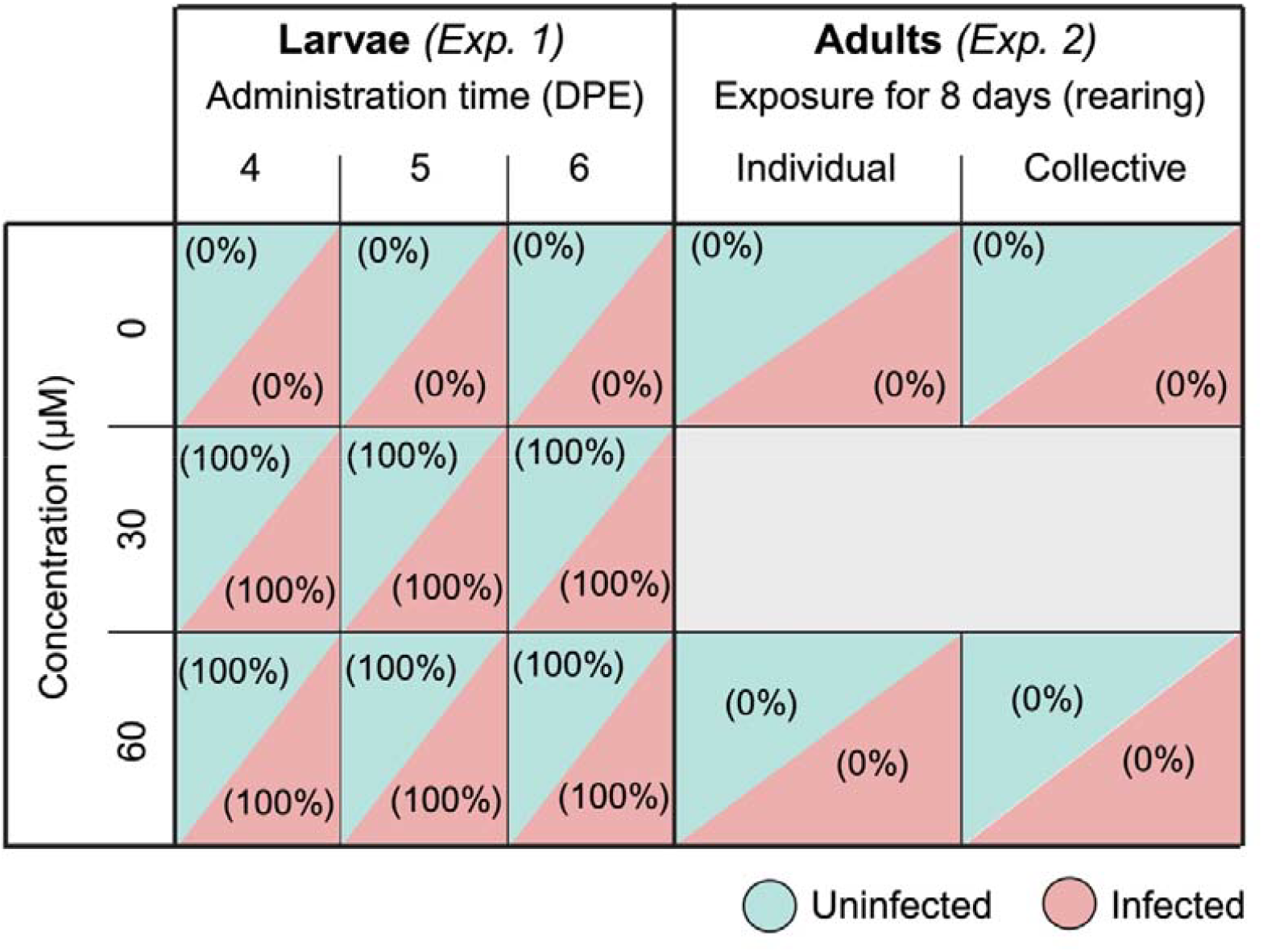
Illustration of the experimental design. Dexrazoxane’s effect on spore load and mortality was tested on both larval **(a)** and adult stages **(b)**. In the first experiment, one of three doses (0 µM, 30 µM, or 60 µM) was administered at one of three time-points (4-, 5-, or 6-days post eclosion (DPE)) to infected and uninfected larvae, in a fully combinatorial manner. In the second experiment, we tested for the effect of exposure to dexrazoxane during the adult stage. For this, we supplemented their sugar diet with 60 µM of Dexrazoxane, or not, and tested for its impact on the spore load when reared individually or collectively, and infected or not. Values between brackets represent mortality for that treatment. For more details, see Materials and Methods (section 2).

#### 2.2 Adult exposure to Dexrazoxane

In a second experiment **(Fig. 1)**, we assessed the impact of Dexrazoxane on: i) spore growth at the adult stage, and ii) if being reared individually (Individual) as opposed to collectively could influence this growth (Collective). The latter is to test if Dexrazoxane needs to be administered individually or if it has the same effect when provided to a group of mosquitoes (with higher stress due to higher density). Two-day-old larvae were exposed to either 0 or 10,000 spores of *V. culicis*. Pupae were transferred to individual 50 ml falcon tubes with approximately 10 ml of deionised water (Dao et al., 2010). Female adults were then transferred to different containers based on their rearing treatment: those reared individually were placed into separate 150 ml cups (5.5 cm Ø x 10 cm) covered with a net, while those from the collective rearing treatment were housed together in a cage (21 cm x 21 cm x 21 cm). The cups contained a 50 mm Ø petri dish (to prevent mosquitoes from drowning) floating on the surface of 50 ml deionised water and a 10 × 7 cm rectangular filter paper to maintain humidity. The mosquitoes were provided with cotton balls soaked in a 6% sucrose solution or a 6% sucrose solution supplemented with 60 µM of Dexrazoxane, which was replaced every other day. We used only one dose due to technical constraints. Unlike the previous experiment on larvae, we allowed treated adult mosquitoes to feed on the Dexrazoxane-sugar-soaked cotton for eight days. The reasoning behind this decision was that while at the larval stage, they are constantly ingesting and exposed to Dexrazoxane, in the adult stage, the exposure is not constant but dependent on the mosquito’s motivation to feed and, therefore, extending the exposure time will maximize the ingestion of Dexrazoxane by the adult mosquitoes.

### 3. Statistical analysis

All analyses were conducted with the R software (Team, 2020) version 4.4.1, using the packages DHARMa (Hartig and Hartig, 2017), car (Fox et al., 2012), glmmTMB (Brooks et al., 2017), emmeans (Lenth et al., 2019) and multcomp (Hothorn et al., 2016). Significance was assessed with the “Anova” function of the “car” package (Fox et al., 2012). We used a type III ANOVA in the case of a significant interaction and a type II ANOVA otherwise. When relevant, we performed post-hoc multiple comparisons with the package “emmeans”, using the default Tukey adjustment.

The presence of spores was coded as a binary response (presence or absence of detectable spores) and analysed with a generalised linear model with a binomial distribution of errors, where the explanatory variables were diet (with or without Dexrazoxane), type or rearing (individual or collective) and their interaction. Because we counted the number of spores in a haemocytometer containing 0.1 µl of the sample (i.e. 1/1000 of the total volume), the detection threshold was estimated to be 1000 spores.

Spore load of mosquitoes with positive spore counts was analysed with a generalised linear model with a negative binomial distribution of errors. The explanatory variables were diet (with or without Dexrazoxane), type or rearing (individual or collective), and their interaction.

## Results

Larval exposure to Dexrazoxane delayed larval development and blocked pupation, independently of the dose, with none of the treated larvae surviving past pupation (as opposed to Dexrazoxane-unexposed larvae). That led us to discard the use of this chemical during the larval stage.

The proportion of individuals harbouring spores was lower for the individual treatment than for the collective (0.65 vs 0.85, χ^2^ = 12.25, df = 1, p < 0.001, **Fig. 2a**) but was not affected by the consumption of Dexrazoxane (0.75 vs 0.75, χ^2^ = 0.00, df = 1, p = 0.945) or their interaction (χ^2^ = 0.28, df = 1, p = 0.595). In contrast, the spore load of mosquitoes with positive counts was reduced by approximately 59% in individuals consuming Dexrazoxane, but only when reared individually (4.9 × 10^4^ vs 1.2 × 10^5^ in individual rearing, and 9.5 × 10^4^ vs 6.7 × 10^4^ in collective rearing, interaction type of rearing * diet: χ^2^ = 10.35, df = 1, p = 0.001, **Fig. 2b**). For further details on the statistical analysis see **Table 1**.

**Table 1.** Effect of exposure to Dexrazoxane on infection. Statistical models carried on to test for effect of exposure to Dexrazoxane, rearing condition (individual in cup, or collective in cage) and their interaction on the presence of spores (Model 1a) and number of spores (Model 1b). See Materials and methods for more details on the experimental design and statistical tests. Values in bold are statistically significant.

| Tested effect |  |  |  |
| --- | --- | --- | --- |
| Model 1a – Presence of spores | df | $\chi^2$ | p |
| Exposure to Dexrazoxane | 1 | 0.00 | 0.945 |
| Rearing (Individual/Collective) | 1 | 12.25 | **<0.001** |
| Exposure to Dexrazoxane: Rearing (Individual/Collective) | 1 | 0.28 | 0.595 |
| Model 1b – Spore load | df | $\chi^2$ | p |
| Exposure to Dexrazoxane | 1 | 2.13 | 0.144 |
| Rearing (Individual/Collective) | 1 | 4.68 | **0.030** |
| Exposure to Dexrazoxane: Rearing (Individual/Collective) | 1 | 10.35 | **0.001** |

**Figure 2.**
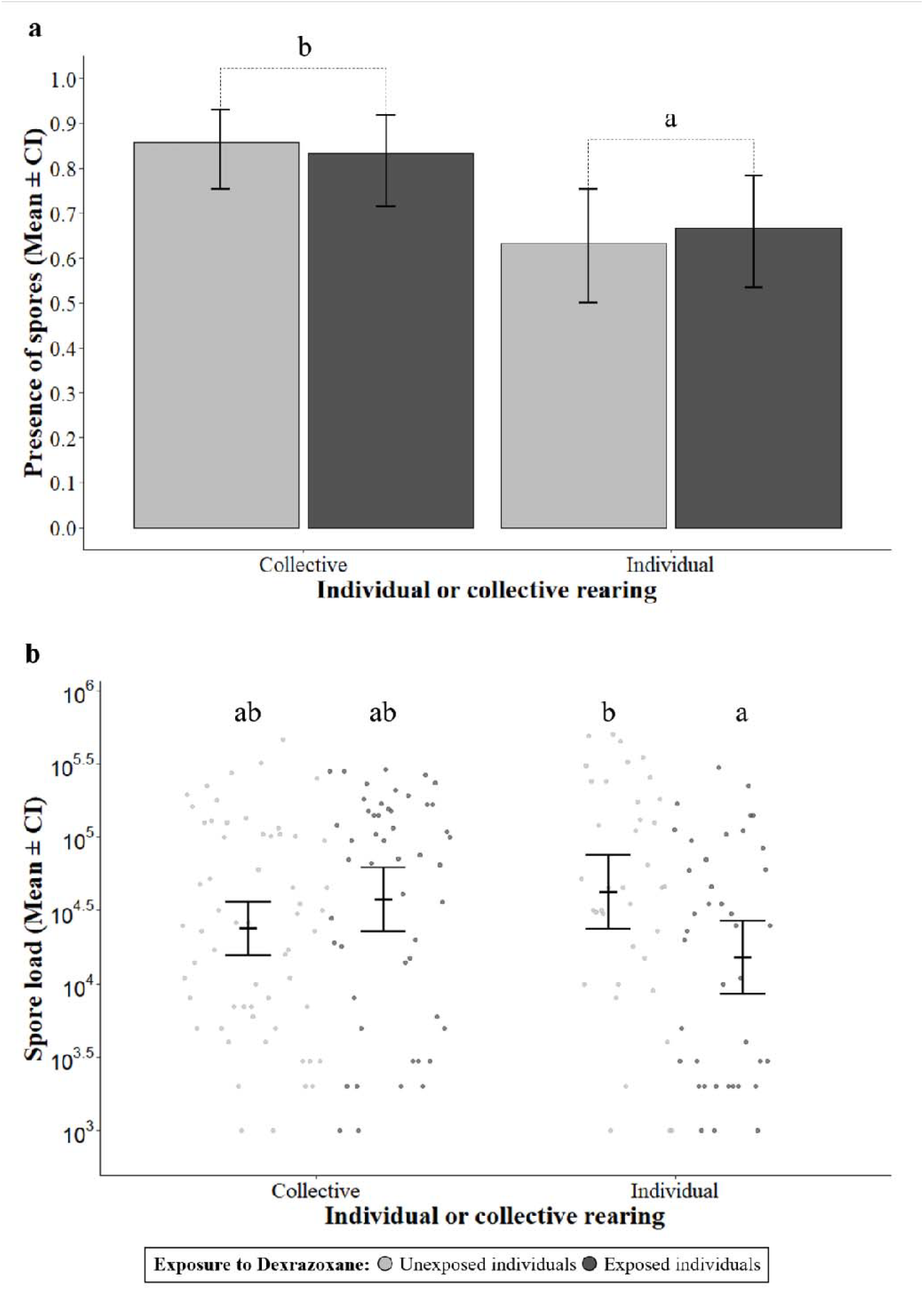
Parasite load of infected mosquitoes after consuming a specific diet for 7 days. The proportion of mosquitoes reared individually or collectively that had detectable spores for **(a)** each Dexrazoxane diet, and their spore load **(b)**. The sample sizes were 70, 60, 60 and 60 for **(a)**, and 60, 50, 38 and 40 for **(b)**. Error bars show the 95% confidence intervals of the means. Letters indicate statistically significant differences from multiple comparisons. See Table 1 for further details on the statistical analyses.

## Discussion

Our study investigates the effect of Dexrazoxane on the infection by the microsporidian *V. culicis* in the mosquito host *An. gambiae*. The findings extend the current knowledge of microsporidia-mosquito interactions and suggest potential strategies for controlling microsporidia infections and mosquito-borne diseases like malaria.

The primary finding from this study is that Dexrazoxane significantly reduces the spore load of infected mosquitoes but only under certain rearing conditions, mainly when mosquitoes are raised individually during the adult stage **(Fig. 2b)**. Moreover, although the total number of spores produced decreased, the number of successful infections remained unchanged compared to unexposed individuals **(Fig. 2a)**. This finding is consistent with previous research in *C. elegans* (Murareanu et al., 2022), which reported that Dexrazoxane inhibits microsporidian development primarily by reducing spore production. However, further studies are needed to determine whether Dexrazoxane acts by limiting the initial integration of spores into the host gut wall or by suppressing subsequent spore proliferation, as both mechanisms could account for the observed reduction in spore load. Importantly, this decline in spore numbers did not come at the cost of host survival—100% of the mosquitoes exposed to Dexrazoxane survived the entire duration of the experiment and were sacrificed only at its conclusion for spore quantification.

For future experimental purposes, we tested for the effect of the Dexrazoxane when administrated individually or collectively. Collective administration of Dexrazoxane has two main advantages: it requires less Dexrazoxane (as administration still happens through supplemented-sugar-cottons) and allows for experiments to be conducted in cages, which often can ease their feasibility. Our results show that Dexrazoxane works at these concentrations but is only administered individually. Nevertheless, we cannot discard the fact that (for instance) higher concentrations of Dexrazoxane administered collectively could lead to similar outcomes.

The reduced spore load in mosquitoes raised individually in cups highlights a potential indirect effect of Dexrazoxane on the mosquito immune response or microsporidian metabolism. While any hypothesis cannot be ruled out, this inhibition is likely due to the drug’s mechanism of action. For human purposes, Dexrazoxane primarily functions through iron chelation, where it blocks circulating iron from being used to fuel oxidative stress and tissue damage. Although this mechanism has been shown to affect *Plasmodium falciparium* development *in vitro* (Loyevsky et al., 1999), the same cannot be said for microsporidia. In fact, *C. elegans* infection with the microsporidia *N. parisii* can be blocked using Dexrazoxane but cannot be reverted through iron supplementation (Murareanu et al., 2022). This result suggests inhibition in microsporidian infections might not happen via iron chelation but instead through another mechanism of action. Interestingly, Dexrazoxane also acts as a DNA topoisomerase II inhibitor, which likely explains why larvae exposed to the drug failed to emerge as adults and had their development arrested at the pupa stage. Given the mechanism of action of Dexrazoxane is fair to assume that this DNA topoisomerase II might be particularly relevant for metamorphosis and transition to the adult stage. In agreement with this, a study on *Ae. aegypti*, larvae treated with etoposide (another DNA topoisomerase II inhibitor) exhibited morphological defects and low survival during both larval and pupal stages (Santos et al., 2021), potentially pointing to a similar result. Therefore, it is plausible that *V. culicis* relies on the same protein or a structurally similar one, and inhibiting this protein during the parasite growth phase might block spore production and infection progression. Although intuitively, this positions Dexrazoxane as a possible larvicide agent, there are a couple of reasons why it might not be the best solution to control mosquito populations. First, our study and others have demonstrated that Dexrazoxane might act at the most basic cellular mechanisms that most animals share. Therefore, there is a long-term potential for environmental damage if this drug is used to target the mosquito larval stage. The use of Dexrazoxane in water bodies where mosquitoes live is likely to impede the correct development of other arthropods or even fish that share the environment. Second, it is crucial to consider that in these experiments, Dexrazoxane was used at relatively high concentrations but in small volumes of water. In the wild, water bodies would have much higher volumes of water, which would raise the cost of the control strategy.

It is important to note that the context of mosquito hosts introduces additional complexity compared to previous studies performed *in vitro* or with *C. elegans* (Loyevsky et al., 1999; Murareanu et al., 2022). Dexrazoxane’s reduction in spore load, especially in isolated conditions, suggests that other factors, such as social interactions and spore transmission, may influence its efficacy. These factors should be carefully controlled and studied in future research, along with determining the optimal dosage. Despite these complexities, the results of this study open the door for further investigation into microsporidia as potential biological control agents, particularly for *V. culicis*. Moreover, the strategic timing of the suppression of the infection progression, without completely clearing it, offers several advantages for studying interactions with vector-borne parasites, such as *Plasmodium*. We believe this approach will allow researchers to investigate the mechanisms behind microsporidia-driven inhibition, for instance, whether it is driven by collateral immunity or resource competition. A further understanding of these mechanisms will allow us to better prepare and design vector control strategies, which are essential today.

## Acknowledgements

We thank Gwendoline Acerbi for her technical support and Jacob C. Koella for early feedback on the project. LMS and TGZ were supported by SNF grant 310030_192786.

## Author contributions

LMS and TGZ designed and conducted the experiments, analysed the data and wrote the manuscript.

## Data availability

All data generated or analysed during this study are included as Supplementary Information files.

## Additional Information

The authors declare no competing interests.

